## Supplementary material for "Lipids modulate the organization of VDAC1, the gatekeeper of mitochondria": Sfigues

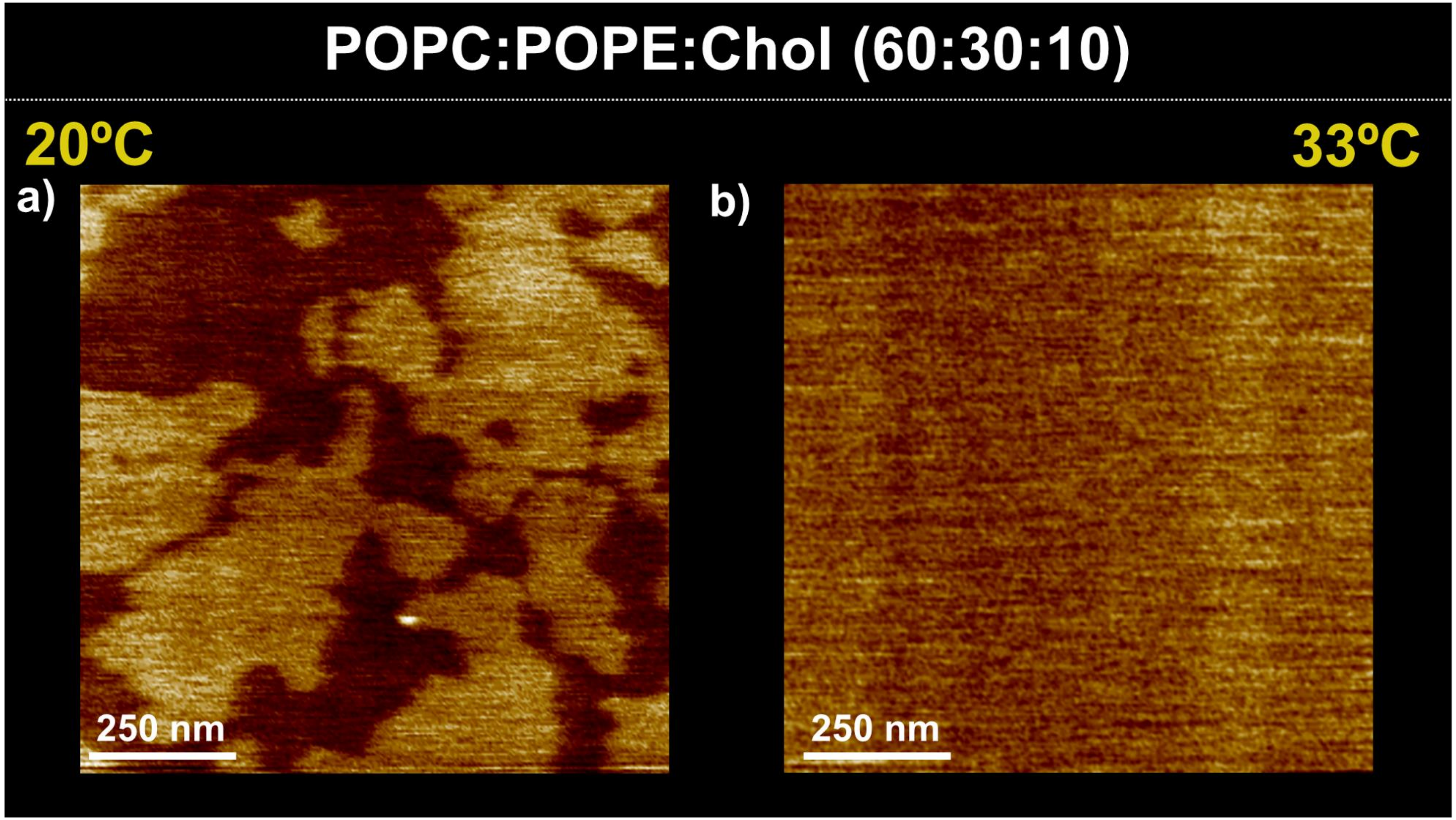


**SFig 1: Cholesterol induces a phase separation at low temperature.** AFM topograph of pure lipid membranes of POPC:POPE:Chol (60:30:10 w:w ratios) at a) 20°C shows a phase separation, and b) at 33°C without phase separation. Similar controls were performed for all conditions, all showing a single phase at 33°C.

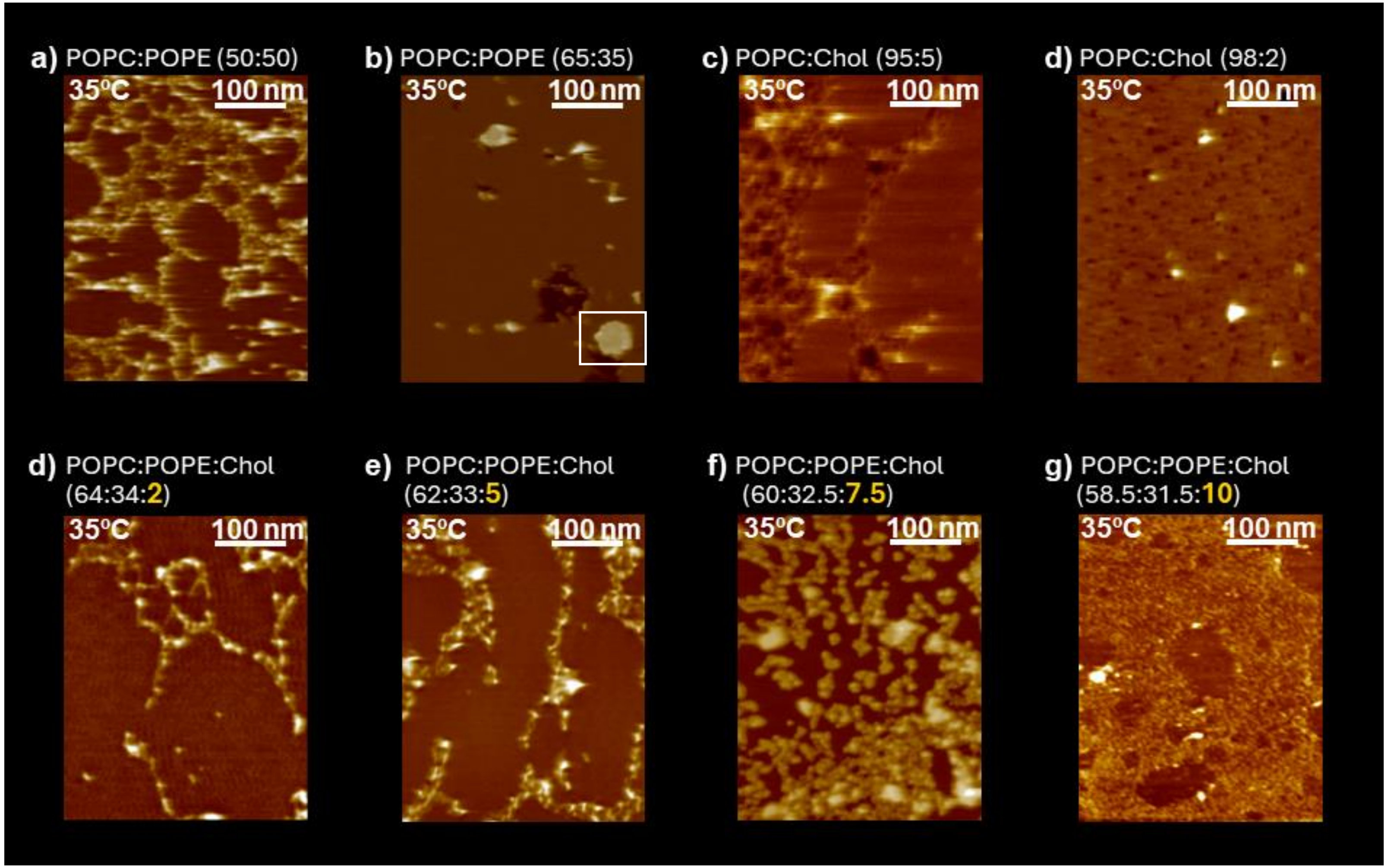


**SFig. 2: VDAC can adopt different assemblies in different lipid compositions.** Gallery of AFM topographs of mVDAC1 in membranes of different lipid composition (w:w ratios). Height false color scale in all images is 11 nm. Each condition is described at higher magnification in following figures. Fig. 2a) is a zoom in the white square in b); a) and c) are described in Fig S6; d-g) are described in Fig. 4.

| Lipid | % weight |
|---|---|
| PA | 1 |
| PC | 54 |
| PE | 29 |
| PS | 2 |
| PI | 13 |
| SM | 0.5 |
| Chol | 9 |
| CDL2 | 0.5 |

**Table S1:** Polar head lipid composition of the mitochondrial outer membrane in weight ratios, based on Daum & Vance 2013. Note that this is a composition from rat liver mitochondria outer membrane. Variation in lipid content of MOM can vary from the tissue, physiological state of the cell or mitochondria function.

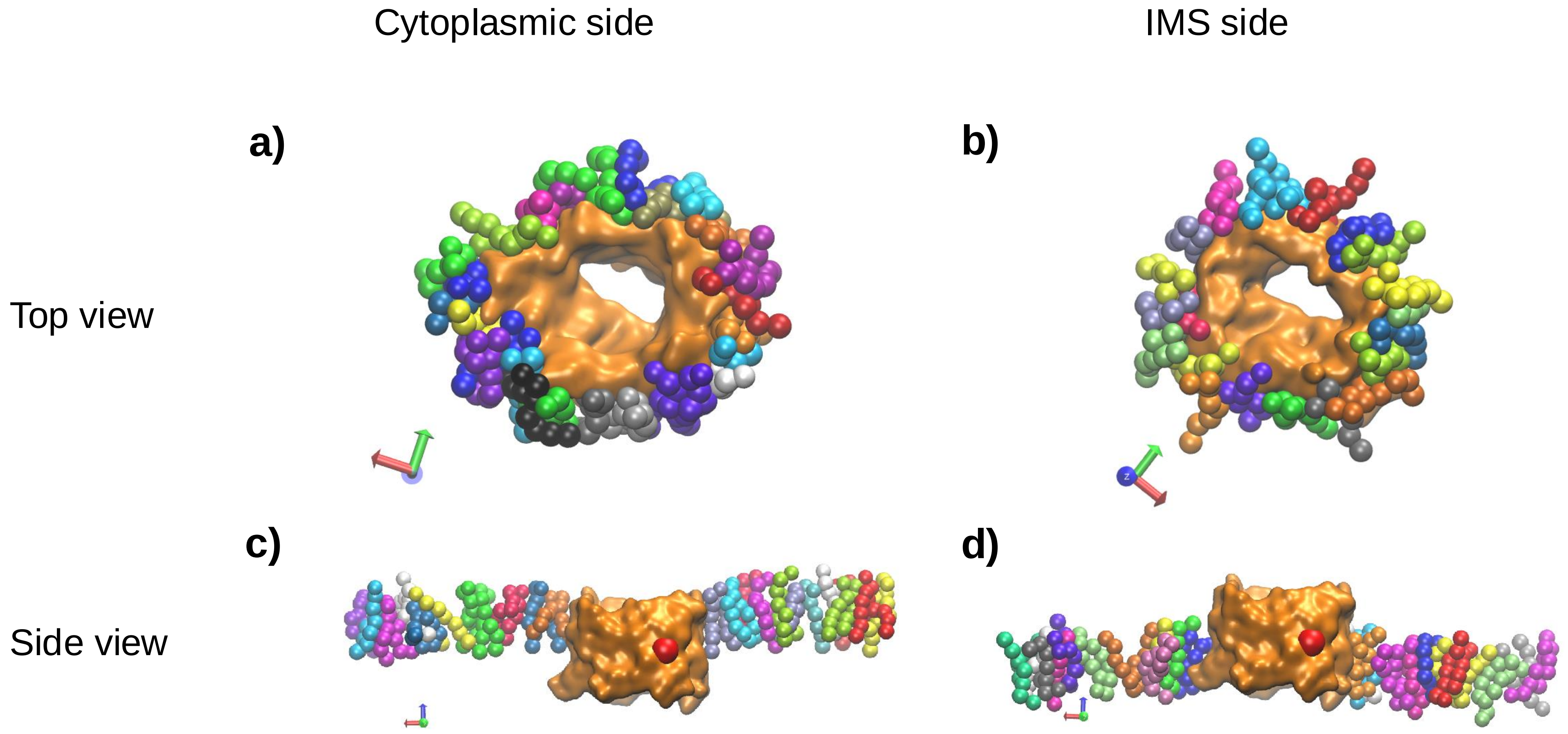


**Sfig. 3: Lipid accommodation on the surfaces of VDAC1**. top (a-b) and side (c-d) views. Approximately twenty lipids are in contact with the protein in the first lipidation shell, either on the cytoplasmic side (a) or periplasmic side (b). Additionally, there are approximately 10 to 13 concentric layers that form the lipid shells around the protein. The color scheme highlights individual lipids. The red sphere on the surface of hVDAC1 represents the Glu 73 side chain, a charged aminoacid facing the membrane.

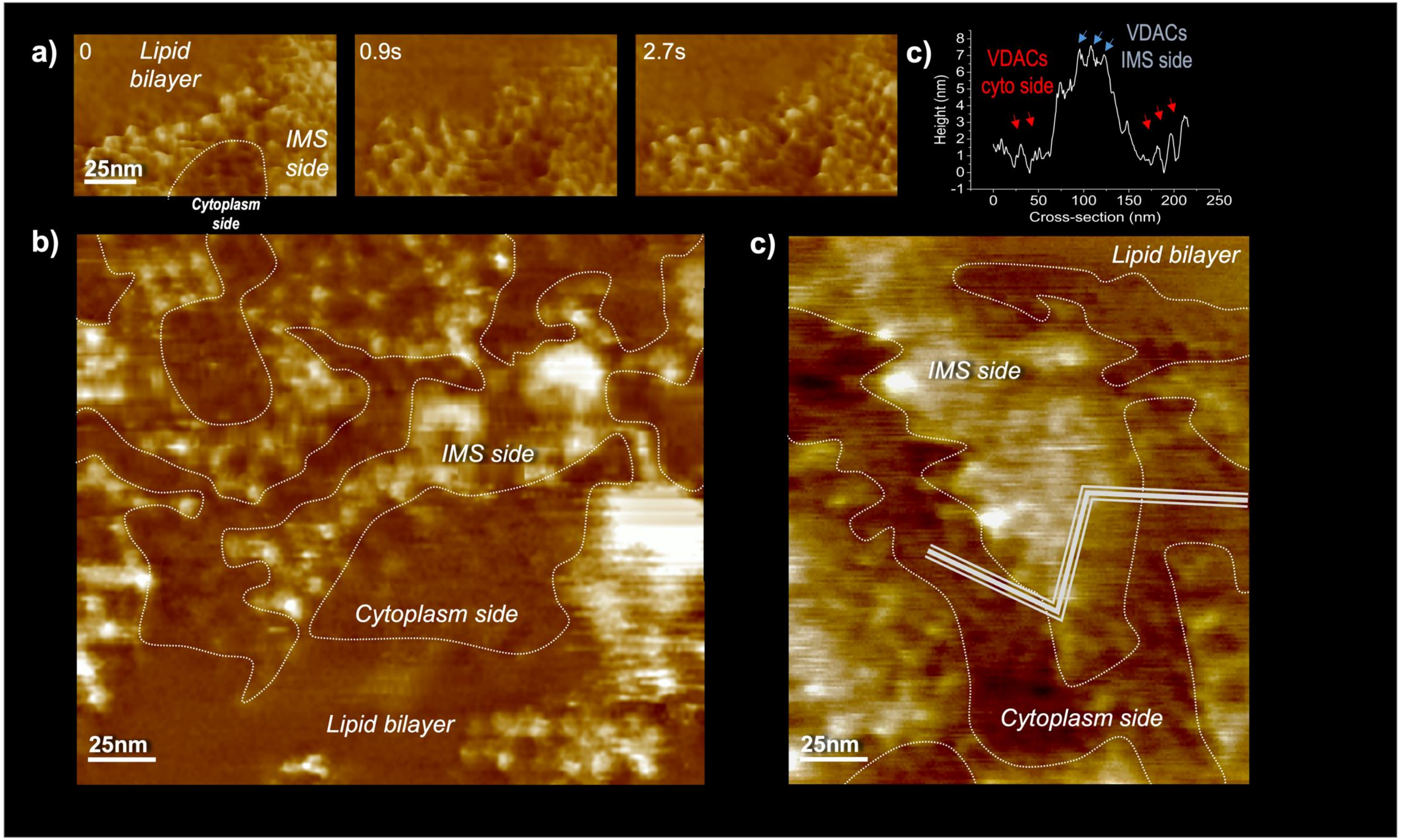


**Fig S4 : AFM topograph show parallel clusters of VDAC in both orientations.** Gold nanoparticles were added to VDAC1 reconstituted in POPC/POPE/Chol (60:32.5:7.5 w:w ratios) and imaged by AFM. a) stability of the gold nanoparticle on VDAC. b) Larger view of both orientations. c) Height profile of a cross-section showing both orientations of VDAC.

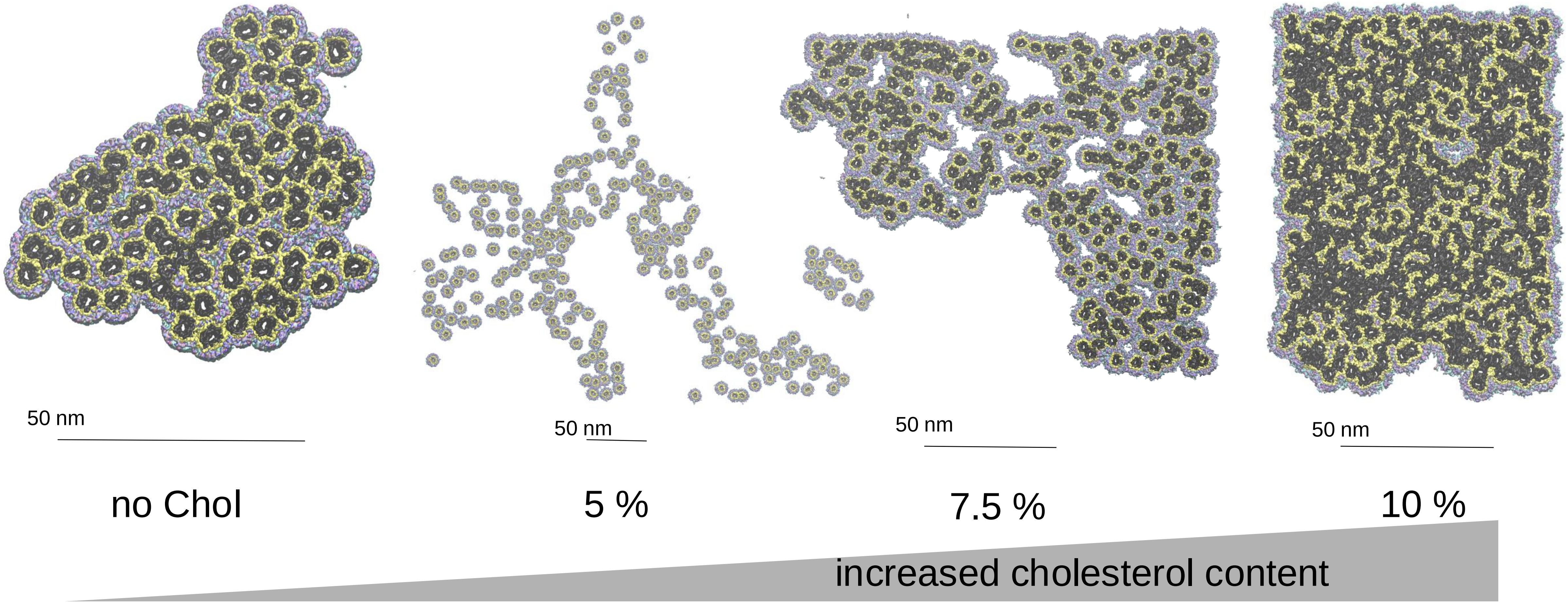


**SFig 5: Cholesterol controls the compaction of VDAC clusters.** Higher magnifications of moleculare models from Fig. 2 and 4. VDACs are in dark grey, bound lipids in yellow and the other interstitial lipids in an atom color-coded representation

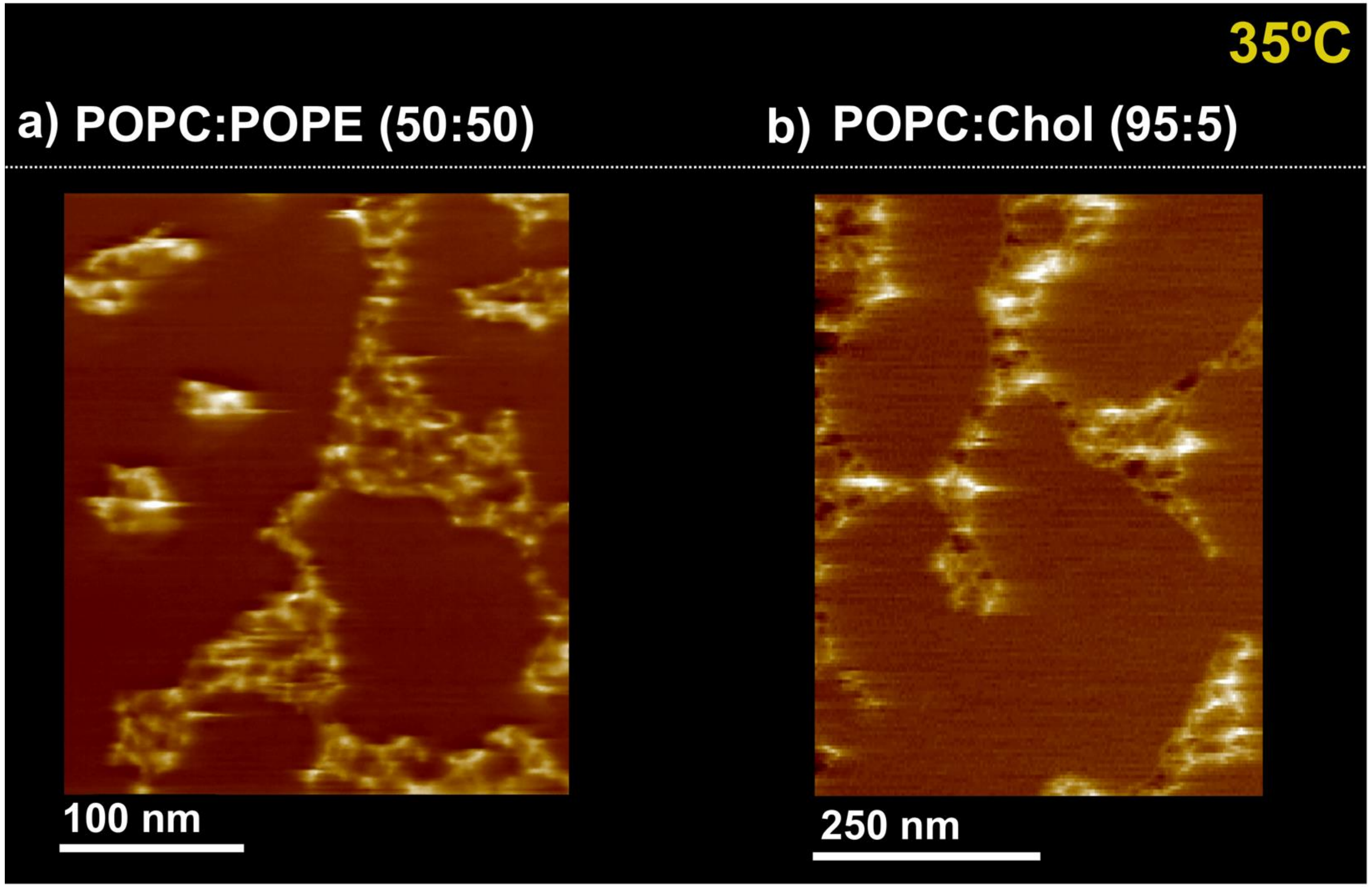


**SFig 6: Lipid ratios outside of the physiological range fail to form glass-like assemblies.** AFM topograph of VDAC in membranes composed of **a)** POPC:POPE (50:50 w:w ratios), or b) POPC:cholesterol (95:5 w:w) show more dynamic structures, different from the honeycomb-like glass-phase clusters observed in Fig. 2 and 4. Height false color scale: 11nm

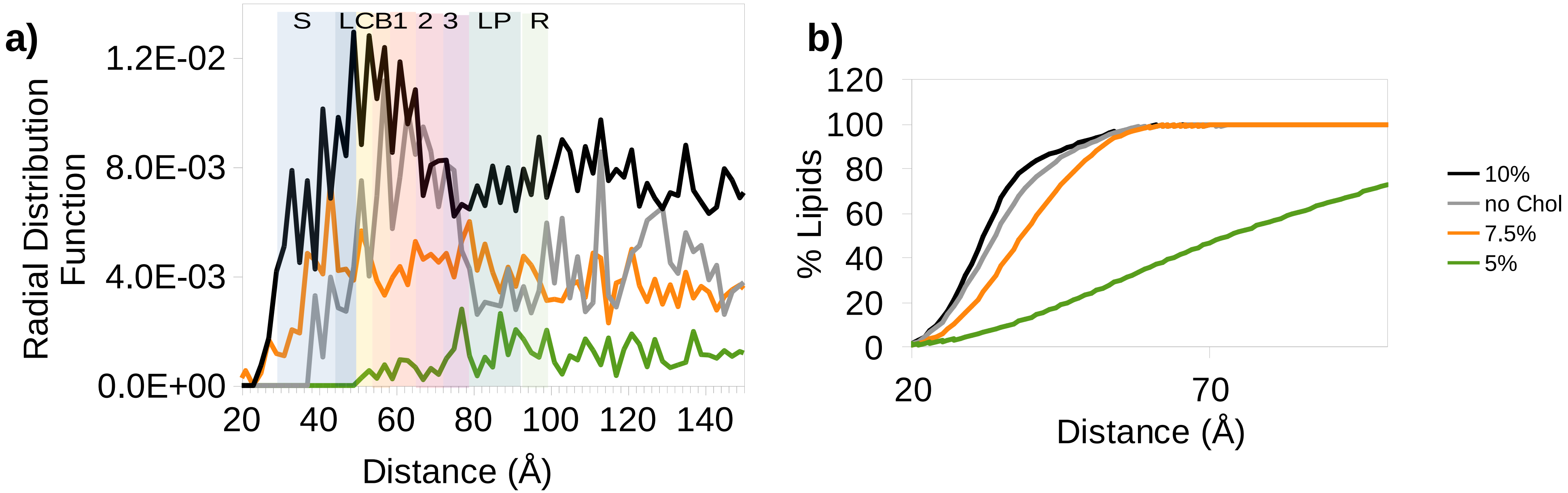


**Sfig. 7: : Proximities between Protein-Protein and Lipid-Protein in the model assemblies**
**a:** VDAC-VDAC 2D Radial Distribution Function computed from manually selected AFM positions within the range of 0 to 150 Å. The letter code and corresponding color strips represent the different categories observed in the models, namely: S for strong clashes, L for low clashes, C for contacts, B for lipid bound, 1 for first extralayer, 2 for second extralipid layer, 3 for third extralipid layer, LP for lipid or protein bridged, and R for row. Note that for the sake of conciseness, the first three categories were grouped as "contacts" in Figure 5.
**b:** Lipid-Protein proximities calculated as the fraction of lipids present at increasing distances around the proteins.

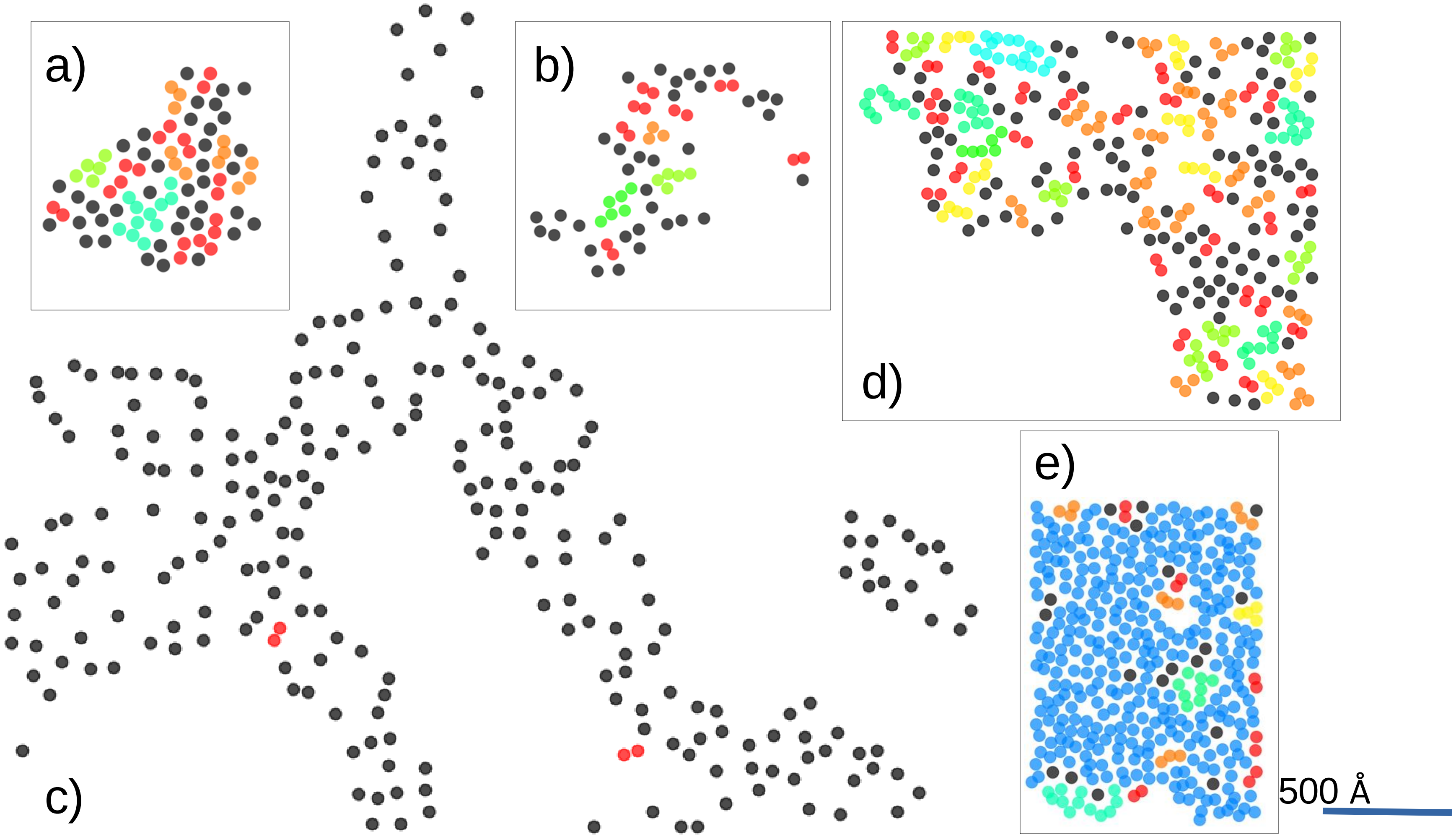


**Sfig. 8: VDAC Clusters in Membrane Assemblies**

This figure illustrates the cluster assignment for VDAC assemblies within a mitochondria outer membrane (MOM)-mimicking membrane, characterized by varying cholesterol concentrations. Panel **a** shows a POPC/POPE (65:35 w:w) membrane, while panels **b**, **c**, **d**, and **e** display the effects of adding 2%, 5%, 7.5%, and 10% cholesterol, respectively. The DBSCAN clustering algorithm utilized a contact distance of 53 Å between proteins, along with AFM-derived 2D coordinates, to assign clusters of varying sizes. Dimers are represented in red, trimers in orange, tetramers in yellow, and higher-order oligomers ranging from five to over 350 are colored from light green to blue. Non-assigned VDAC monomers are depicted in black.

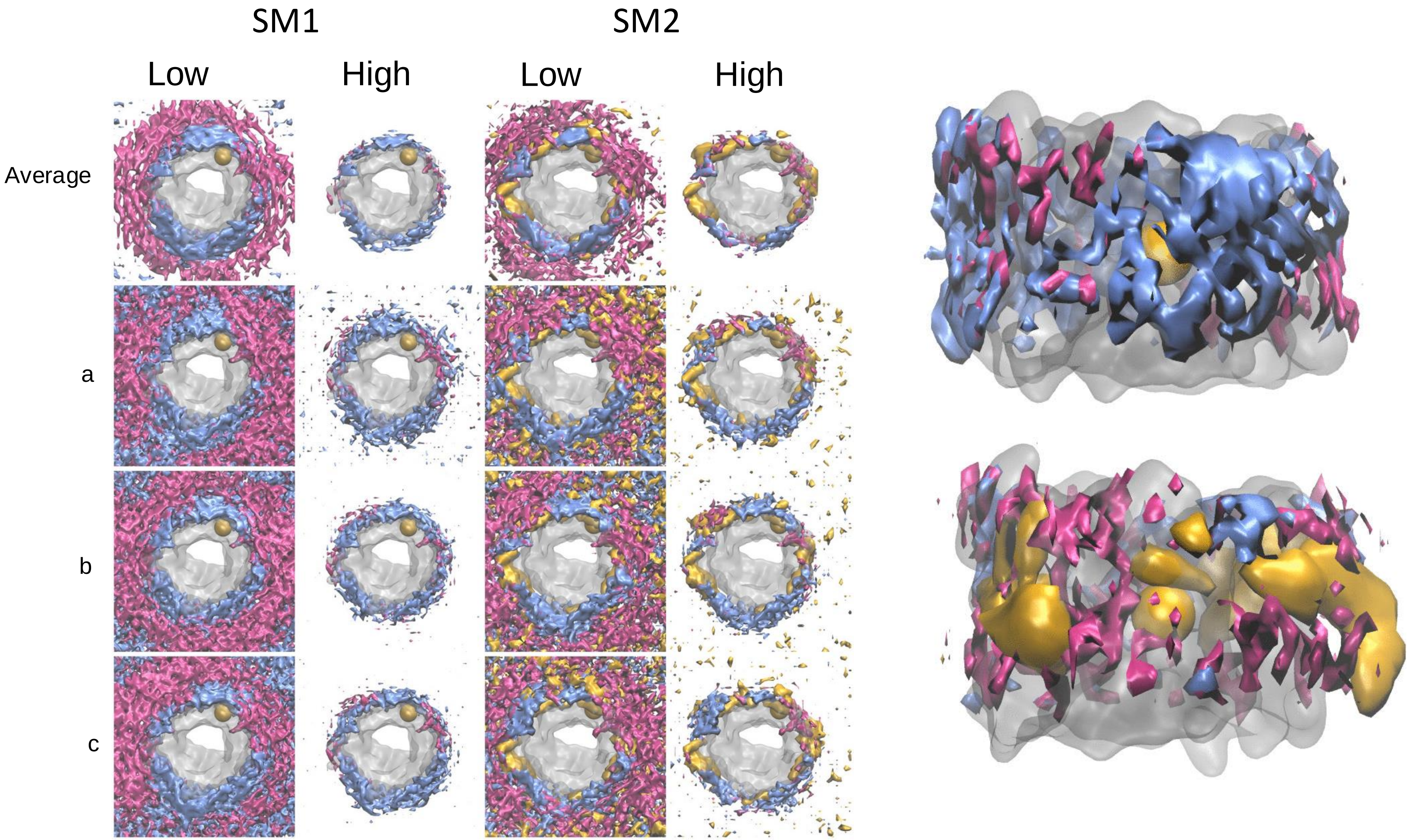


**Sfig. 9: Volumetric maps of lipid occupancies in molecular dynamics simulations of VDAC1**. Left panel : **t**he mean occupancy of POPC (magenta) , POPE (blue) and cholesterol (gold) was calculated over the last 10 µs of 20 µs coarse-grained MD simulations. Three systems (a, b, c) were independently built using the same composition but different distributions of randomly positioned lipids. The upper row shows the averaged map corresponding to the distinct maps shown in the following rows. Low and high densities represent an excess density of 20 and 80%, respectively, relative to the average density.

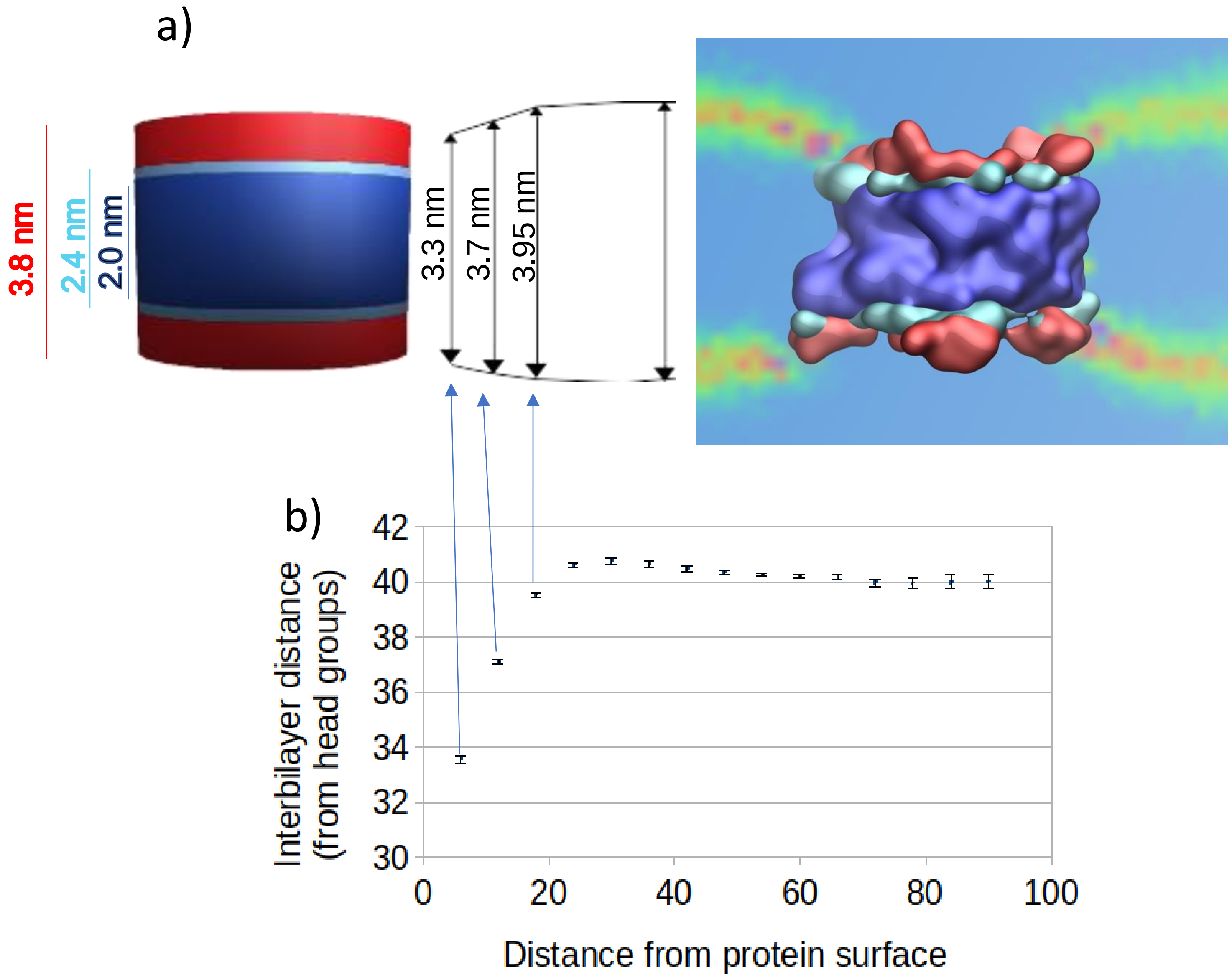


**Sfig. 10: VDAC-to-bilayer adaptation during the last 10 µs MD simulation** in Simple membrane 2 system. **a** The hydrophobic core, an intermediate area, and the hydrophilic extension of VDAC1 are represented using deep blue, cyan, and red, respectively. Thicknesses were evaluated from particle distributions projected on the membrane normal axis using hydrophobic and hydrophilic residues of the protein. The intermediate region corresponds to the area that separates the half-maximum positions of the two distinct peaks. The arrows represent the membrane thickness calculated for the first three successive lipid layers considered at 6, 12, and 18 Å from the protein surface. The last arrow is the thickness measured at a longer distance. Thicknesses were computed from the distribution of lipid head groups, including phosphate, glycerol, and cholesterol OH particles. **b** the evolution of the membrane thickness as a function of protein distance was further characterized using a Gaussian fit to the head group distribution. The error bars correspond to the error estimation on the fit.

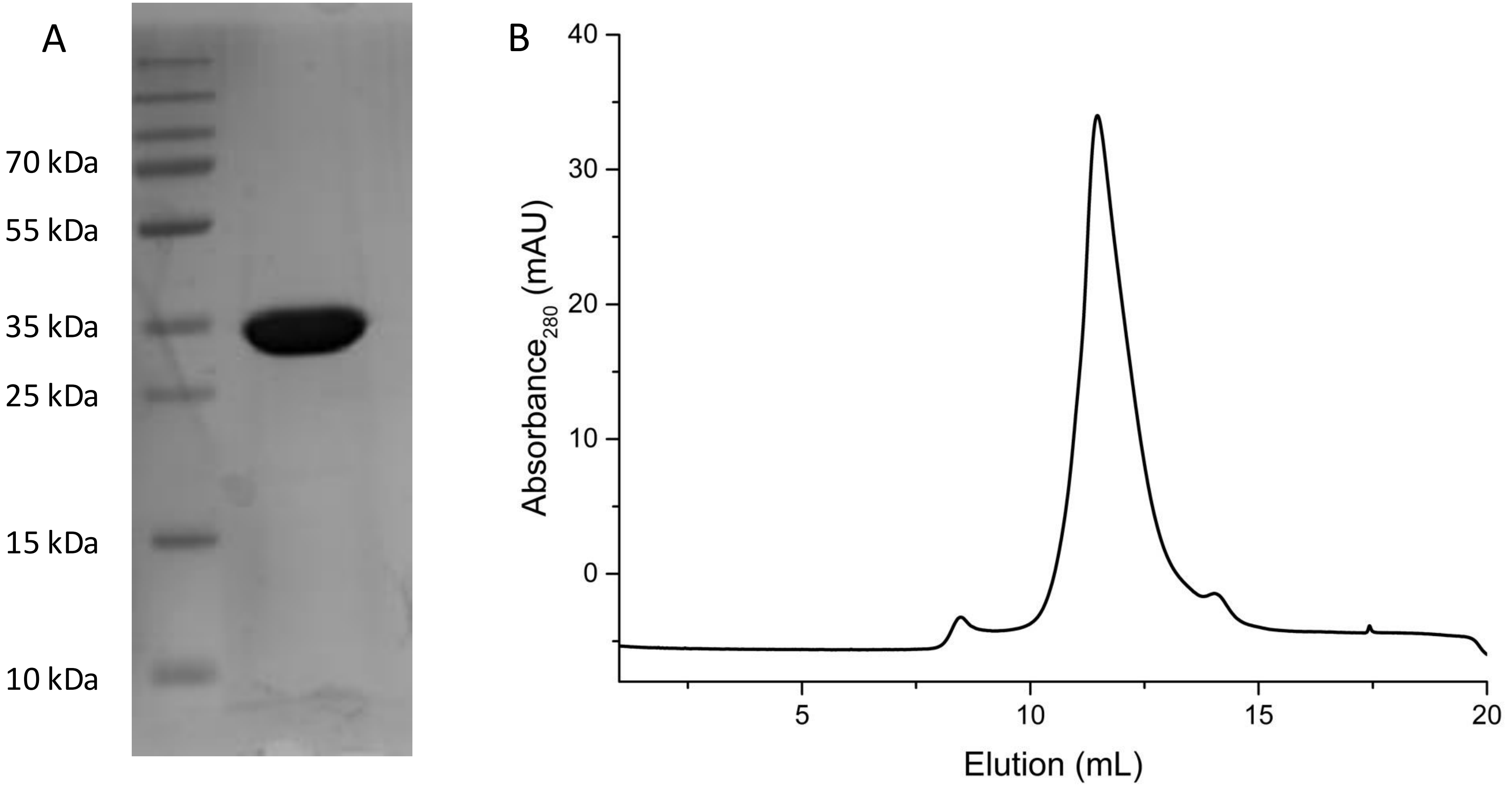


**Sfig S11 : mVDAC1 purification.** A: SDS-PAGE and B. size exclusion profile (superdex 200 10/300 GL) of purified mVDAC1.

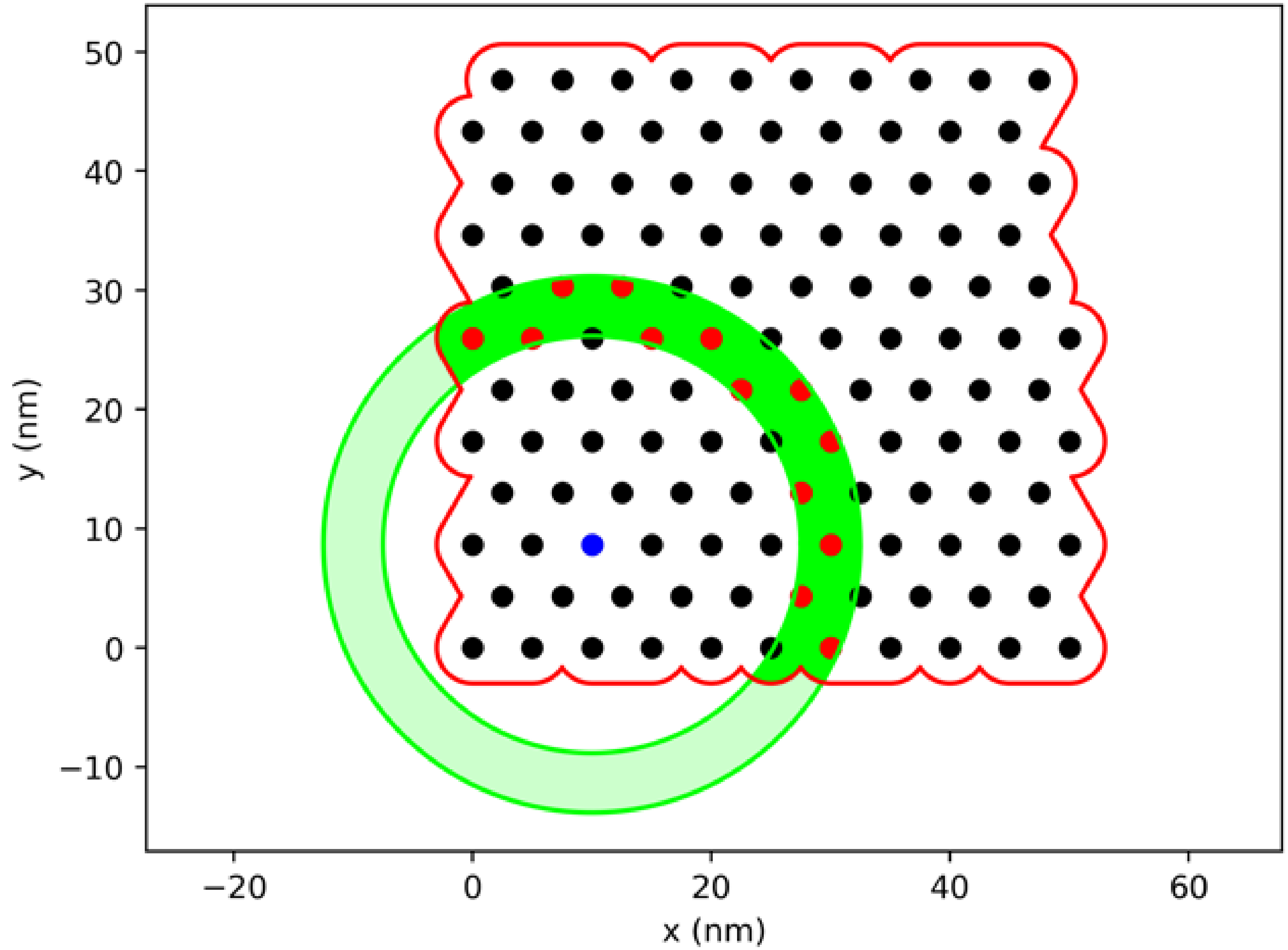


**Sfig S12 : Schematic representation of the RDF calculation**. To caclulate the RDF of molecules within a protein patch first the borders of the patch were determined (red line) and its area (total area) and then for each (blue spot) of the "N" protein molecules within the patch a series of concentric rings were constructed at different distances (r) with a width of dr. Then the number of proteins found in the ring ("n_R", red) were counted as well as the total area within the patch ("a_R", green). These values were summed and finally the RDF (g_R) was calculated as the total number observed at a given distance from another molecule divided by the total area observed (sun n_R/ sum a_R) at that distance relative to the total number of molecules divided by the total area (N/A). The areas ("a_R" and "A") were calculated by monte-carlo integration.

**Supplementary Movie**

HS-AFM movie of a VDAC1 PC PE 2%Chol membrane absorbed on the mica substrate at 33ºC. Movie parameters: frame rate 969 ms; full image of 200 nm x 200 nm and 256x256 pixels; colour depth 8bit (256 values); full colour scale
